# Using cost-effective surveys from platforms of opportunity to assess cetacean occurrence patterns for marine park management in the heart of the Coral Triangle

**DOI:** 10.1101/2020.06.19.160887

**Authors:** Achmad Sahri, Putu Liza Kusuma Mustika, Purwanto, Albertinka J. Murk, Meike Scheidat

## Abstract

The Wakatobi National Park (WNP) at the heart of Coral Triangle is an important area for cetaceans in Indonesia. Currently there is insufficient information on spatio-temporal occurrence patterns of cetaceans to inform effective conservation strategies. This study used platforms of opportunity from May 2004 to May 2012 as a cost-effective way to address this knowledge gap. A database was created of cetacean sightings per surveyed days at sea, allowing for an analysis of species diversity and habitat use around the islands. A total of 11 cetacean species were identified. Spinner and bottlenose dolphins were sighted most often, followed by melon-headed and sperm whales. Spinner dolphin showed a wide distribution in the area, whilst bottlenose dolphin and melon-headed whale occupied the waters between the main islands and south atolls. Sperm whales occurred mostly in waters to the north of the main islands and as melon-headed whales were mostly found in deep waters. Most cetacean sightings occurred in the zones designated for human use, indicating where potential conflicts might occur. No sightings were found in the Park core zone, indicating a mismatch between WNP design and the ecological needs of the cetaceans. A sub-sample of the data from dedicated fishing monitoring trips was used to derive a sighting frequency. Occurrence of both small and large cetaceans was highest during inter-monsoonal seasons, possibly related to an increase of prey availability due to seasonal upwelling and increase in survey activity. Inter-annual occurrence of cetaceans was variable, with no large cetaceans being sighted in 2010-2012, likely due to reduced survey efforts. In areas with limited resources for designated surveys, the use of platforms of opportunity can be a cost-effective tool to provide valuable data on cetacean occurrence. It helps identify potentially important areas as well as highlight where to direct designated research efforts. We discuss the implications of our findings for the conservation management of these cetaceans and give suggestions for improved marine park management.

## 1. INTRODUCTION

Indonesia has a high diversity of cetacean species (Mustika et al., 2015b), and was a popular whaling ground for large whales during the Yankee whaling era from the 18^th^ to early 20^th^ century (Townsend, 1935). The deeper eastern waters of this country are also an important migration route for cetaceans, including large whales (Double et al., 2014). Studies of cetacean species particularly around tropical oceanic islands are limited (Baird et al., 2009), including in Indonesia. These oceanic island cetacean populations likely have specific conservation needs, but these populations are not systematically monitored, particularly in remote areas (Ender et al., 2014) in waters with complex reef and small islands. Some information about cetaceans in Indonesia is available hidden in unpublished internal reports. Insufficient data on spatial ecology of cetaceans impairs effective conservation strategies in this country.

Wakatobi at the heart of Coral Triangle region and part of the Banda Sea ecoregion is the second Indonesian national priority for area conservation (Huffard et al., 2012). Wakatobi is characterized by complex reefs and small-islands with a diverse submerged topography comprising a continental shelf, slopes, and pelagic waters with trough-, ridge- and seamount-like features. These characteristics are known to influence ocean circulation, induce nutrient upwelling and provide a diversity of water masses resulting in a rich habitat complexity, high biodiversity and great abundance for many species including cetaceans (Bartholomew et al., 2000; De Vos et al., 2012). The unique configuration of near-shore yet deep-sea habitat of Wakatobi is of special importance to deep-sea cetacean species, making the area one of the most important marine systems in Indonesia. Wakatobi waters were therefore designated as a marine park in 1996 (WNP Authority, 2008). The near-shore deep-sea cetacean habitats in Wakatobi provides a unique opportunity to survey deep-water cetacean species that normally occur in waters further offshore, which would otherwise be too challenging to monitor with small boats (e.g. in Malaysian waters (Ponnampalam, 2012)). However, there is significant fisheries activity (Pet-Soede and Erdmann, 2003) and the effectiveness of the marine park as a conservation tool for cetaceans is not known.

Unfortunately, as in other regions of Indonesia, data availability on the status of cetaceans in this regionally important habitat (Huffard et al., 2012) is very limited, posing big challenges for cetacean conservation and management. Neither cetacean diversity nor the spatio-temporal occurrence of the different species is known for Wakatobi waters. In addition, the habitat preferences of the cetaceans in this area in terms of seafloor topography such as reef habitat types and depth have not yet been described in detail. This lack of information hampers informed conservation efforts within the area. To address this issue, a cetacean monitoring program was initiated in the Wakatobi National Park (WNP) and adjacent waters in 2004 and cetacean sightings were documented until 2012. The program aimed to assess cetacean species diversity, reveal spatio-temporal occurrence patterns and their habitat type preferences. The initiative for gathering such information was required by the WNP Management Plan document (WNP Authority, 2008).

Identification of areas of particular importance for a species is a key aspect in conservation and management of wildlife. This notion requires the acquisition of baseline information on species distribution and dynamics, which are complex and logistically expensive to obtain for cetacean species via dedicated surveys over such a vast area (Redfern et al., 2006). Ideally, systematic surveys are performed following a sampling design that gives each sampling point in the study area an equal probability of being sampled (e.g. line transect, Buckland et al., 2001). Conducting such an unbiased and elaborate survey generally involves dedicated vessels and observer teams and thus can be prohibitively costly (Williams et al., 2006). An alternative in the case of limited resources has been placing observers on platforms of opportunities, such as inter-island ferries and whale watching vessels (Kiszka et al., 2007; MacLeod et al., 2009; Williams et al., 2006). Assuming these vessels cover similar areas and the observer effort is also comparable over time, this data can be used as an index of occurrence (e.g. sighting frequency). Such long-term datasets constitute a valuable source of information for investigating how cetaceans use an area (Arcangeli et al., 2013; Silva et al., 2014). The occurrence of cetaceans over time and space is a crucial input for designing effective spatial planning, to evaluate park zoning systems and to prioritize measures for protecting cetaceans from adverse human impact.

The threat of adverse human impact especially occurs from spatial and ecological overlap with human activities (Williams et al., 2006). Important human disturbances include direct (bycatch) and indirect (prey depletion) impacts from fisheries (Reeves et al., 2013), as well as physical and acoustic disturbance mainly by marine traffic (Erbe et al., 2019; Pennino et al., 2017), seismic activities from oil and gas exploration and naval sonars (Henderson et al., 2014; Rosenbaum and Collins, 2006), and various sources of pollution (Allen et al., 2011; Tanabe, 2002; Venn-Watson et al., 2015). Increasing anthropogenic stressors include coastal-offshore development and energy production, resource extraction, tourism, and climate change (MacLeod, 2009; Passadore et al., 2018b). To assess the potential impact from these human threats, baseline information on cetacean occurrence at a local level is urgently needed.

Our objective was to develop an approach to use platforms of opportunity as well as incidental data from long-term visual monitoring to add knowledge of unstudied cetaceans populating oceanic island-based habitats of Wakatobi. This approach provided information on the diversity, spatio-temporal occurrence patterns, and relative abundance of cetaceans in the area of interest. By comparing the spatio-temporal occurrence patterns with bathymetric characteristics and reef habitat types, we also revealed habitat preferences of these cetaceans. Finally, we discussed the advantages and caveats of using non-systematically collected data, and the implications of our approach and findings for effective cetacean conservation, spatial planning and marine park management.

## 2. MATERIALS AND METHODS

### 2.1. Study area

The study area encompasses Wakatobi National Park (WNP) and adjacent waters, centrally located within the Coral Triangle region, a region with exceptional marine biodiversity (Green and Mous, 2008). The park includes a remote island group, approximately 120 km off the southeast Sulawesi mainland (Figure 1). The area is one of the largest marine parks in Indonesia and covers approximately 13,900 km^2^, containing all the major reef types. WNP includes four major islands with the boundaries are congruent with those of the Wakatobi district government (Clifton, 2013). The area has a diverse and complex submerged topography with channels between major landmasses that serve as passages for migrating mammals (Pet-Soede and Erdmann, 2003). The area has a relatively narrow continental shelf and the depth increases very rapidly from the shelf edge at around 150 m from the shorelines. The oceanographic conditions are influenced mainly by circulating and seasonally changing currents in the Flores and Banda Seas and result in productive and relatively cool waters as a consequence of upwelling from the south (Pet-Soede and Erdmann, 2003).

**Figure 1.**
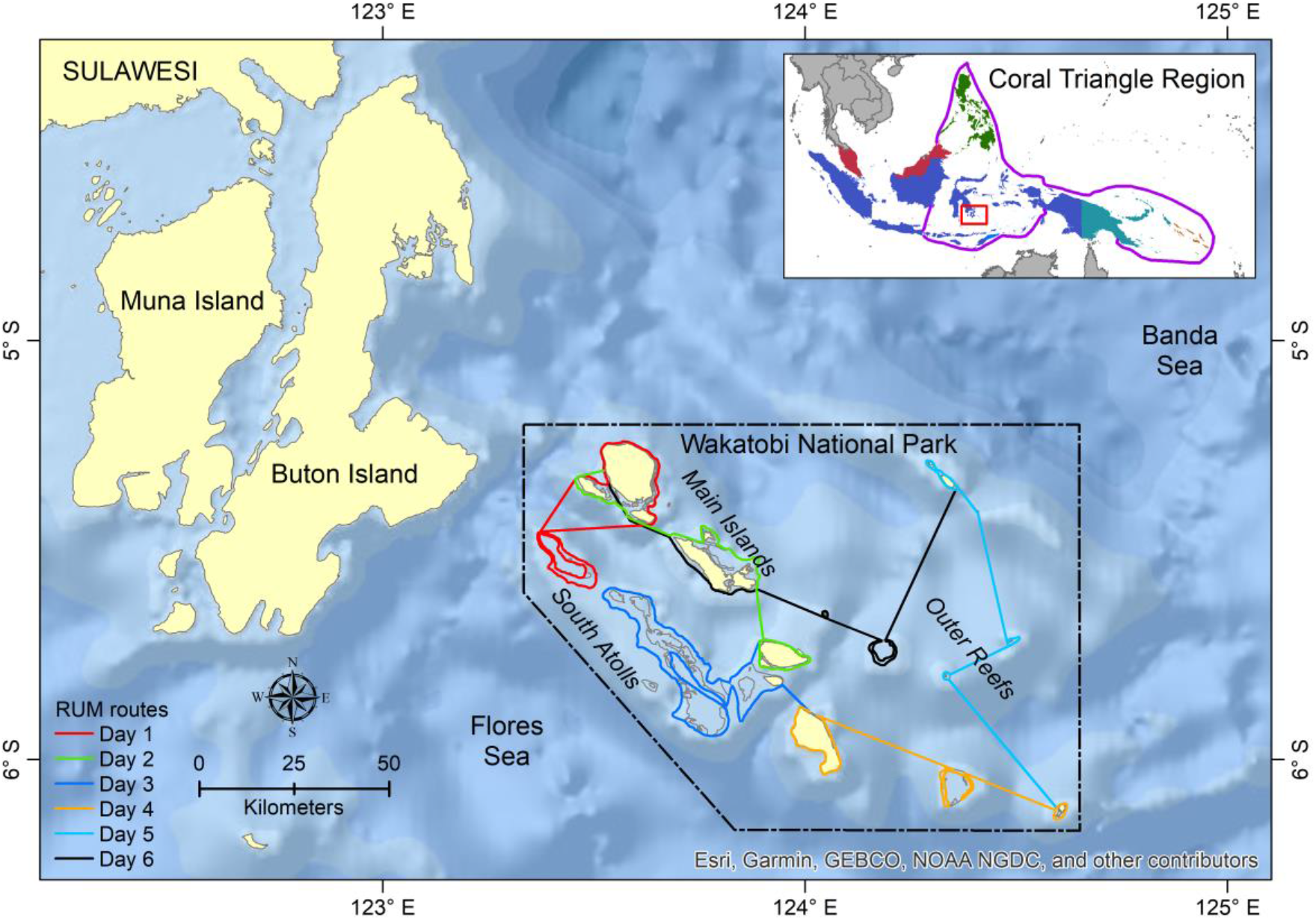
Study area in Wakatobi National Park (WNP) and adjacent waters. The routes of Resource Use Monitoring (RUM) as a main platform-of-opportunity survey are presented as coloured lines.

### 2.2. Data collection

Cetacean data was collected from May 2004 to May 2012 following a cetacean monitoring protocol developed by the WNP Authority and The Nature Conservancy-World Wildlife Fund for Nature (TNC-WWF) Joint Program Wakatobi. Briefly, the protocol was applied by trained personnel mainly during Resource Use Monitoring (RUM) trips conducted to collect data on fishing activity. Other platforms of opportunity used were vessels conducting diverse monitoring programs (reef health, turtle, seabird, spawning aggregation site), the tuna tagging program, VIP guest trips, trips to outer Wakatobi, incidental patrols, and inter-island cruises; thus making the cetacean surveys cost-effective. Sightings reported by fishermen and volunteers were included as incidental sightings.

Different fleets were used as observation platforms, including some speedboats (length ~4 m; height 0.5-1 m; speed ~20 knots); and ‘Floating Ranger Station’ liveaboards (length ~20 m; height 3-5 m; speed 7-9 knots). Travel speed was not constant depending on navigation, boat type and weather conditions.

The RUM was designed to cover all habitat types within the WNP (i.e. main islands, south atolls, and outer reefs), and took six days to complete by following fixed line transects (Figure 1). However, the boats did not always follow the predetermined survey routes due to technical issues e.g. logistics and supply, vessel reparation, and weather constraints. This arrangement resulted in changes to the fixed routes and difference in area coverage between surveys. In 2007 and 2008, several surveys could be conducted in one month, resulting in the higher sum of surveyed days per month (a complete trip suggested by the protocol).

Cetacean observations took place en-route between the harbor of origin and the destination (e.g. fishing locations, location of targeted monitoring) as well as en-route between destinations. Search efforts were suspended during the RUM interview stops or when a targeted monitoring had to take place. At least one observer maintained visual watch during daylight hours (06:00–18:00h, weather permitting). Searching was primarily done with the naked eye or sometimes with the aid of hand-held binoculars. The observers were trained and experienced in cetacean data collection and identification, although their abilities varied and it cannot be guaranteed that search effort was continuous in all cases. All records were quality controlled by experienced observers that were onboard to ensure that the methodology was as consistent as possible between observers. Surveys also recorded days at sea with no sightings. Because the method does not follow a standardized distance sampling protocol (Buckland et al., 2001), all sightings recorded during these surveys were categorized as ‘opportunistic’.

When a cetacean group was detected, the boat slowed down or stopped, and information was recorded on a standardized form. This form included entries for geographic position, time of sighting, species identity, number of individuals, and observer name(s). In the form, estimated distance and relative angle from the vessel to the cetacean group at the time of the sighting were noted as well as specific behavior and other remarks, although not all observers recorded all this information. A cetacean handbook was used to aid in the identification of sightings to the species level. When species identification was questionable, photographs were shown to other cetacean experts working in the region. If identification was not possible on a species level, the sighting was recorded as dolphins, whales or unidentified cetacean. Unidentified species can be caused by a combination of factors, such as very short encounters, distance of the sighting, no clear appearance of the cetacean, or lack of observer identification skills. Weather conditions were not consistently recorded each day and records were not always provided with a sea state description (e.g. Beaufort scale), although outstanding sea conditions including rain, strong wind, and high waves were usually recorded. Vessels did not go out when weather condition were so bad (approximately Beaufort >4) that they would hamper the vessels to conduct their work.

### 2.3. Data analysis

#### 2.3.1. Survey efforts

The survey set-up did not facilitate the collection and storage of continuous GPS data, so information on km tracks or search hours per day were not recorded. The best available proxy for survey effort was survey days at sea. For each survey day, information on cetacean sightings, including days with no sightings, were available. The limitations of the data in terms of survey coverage and number of sightings meant it was not feasible to compare the results from outside and inside WNP. Therefore, we treated our data (inside and outside WNP) as one dataset. All data that included effort information was considered to be part of the “eligible survey” dataset.

An eligible survey was defined as any trip that consisted of four (equal to two third of a complete trip suggested by the monitoring protocol) or more days. This subset of the available data made the different surveys more comparable in their area coverage and applicable as a proxy of effort. Using both ‘days with’ as well as ‘days without’ sightings per survey allowed the calculation of a relative sighting frequency (of both sightings and individual animals).

#### 2.3.2. Datasets used

All cetacean sighting records were quality-checked and placed in a database. This information was used to provide a species record for the area and to describe the presence only occurrence for each species.

With this data filtering, the number of surveys for calculating sighting frequency (i.e. number of sightings per survey day) was reduced to 103 (74%) surveys from the original 140 surveys. The number of sightings was also reduced to 241 (67%) sightings from the original 358 sightings. Therefore, we combined the sightings based on higher taxa and defined large cetaceans (baleen and sperm whales) and small cetaceans (the rest taxa). The sighting frequency was averaged per survey and then monthly and annually.

To understand a possible determinant to the inability to identify cetaceans, we conducted logistic regression (Field, 2013) between the dummy variable of unidentified cetaceans and several independent variables, i.e. the distance between the animals and the observers, the number of animals encountered in a group and season. We could not examine the association between other variables (e.g. the encounter period or observers’ skills) due to the absence of such data.

#### 2.3.3. Spatial and temporal occurrence patterns

All sightings that were obtained from all survey days at sea were combined into a single dataset. ArcGIS 10.6.1 (Environmental Systems Research Institute, Inc.) was used to visualize species spatial distribution or occurrence patterns maps based on sighting locations and number of individuals. Sufficient sightings for depicting spatial occurrence patterns were only available for the four most abundant species: spinner dolphin, bottlenose dolphin, melon-headed whale, sperm whale, as well as two unidentified taxa (dolphins and whales). The cetacean occurrence was correlated to the current park zoning system, reef habitat types, and depth preferences to assess area preference. Depth data was extracted from the General Bathymetric Chart of the Ocean (GEBCO^1^). For investigation of the temporal occurrence patterns, sighting data per species was grouped by month from all survey years. The sighting data of the four most abundant species described in this paper was also used to analyze the habitat suitability of these species in another paper using a more complex habitat model.

#### 2.3.4. Additional important information

Information on cetacean behavior, mother-calf pairs, and cetacean-fishing vessel and fish aggregation devices (FADs) interaction was not always recorded for each sighting. Since the amount of additional data was limited, no inferential statistical analysis could be performed, therefore it only was reported descriptively. Behavior of cetaceans was classified into four categories: (i) travelling – normally moving animals on a steady course, (ii) resting – stationary in one place, almost without movement, (iii) socializing – clear and constant interaction between the animals in a normally stationary group, and (iv) foraging – non-synchronized movements and very active animals, normally involving the visualization of prey or aggregation of birds (Alves et al., 2018). We added (v) ‘bow riding’ as a special category, since the behavior is prevalent in small cetaceans, and it shows a cue of being attracted to vessels (Anderwald et al., 2013).

## 3. RESULTS

### 3.1. Efforts and sighting summary

A total of 671 days were surveyed from May 2004 to May 2012 of which 229 days were with sightings (Table 1). The total of 358 sightings corresponds to 12,846 individual animals. Surveyed days at sea were relatively consistent over the years, although only in 2007 and 2008, all months of the year were covered (Table 1, Figures 2 and S1). Although the months in which surveys were performed included all four monsoonal seasons, most effort was concentrated in inter-monsoonal (transition) seasons characterized by calm weather (Figure 2): from March to May (Transition 1 season, before SE monsoon) and September to November (Transition 2 season, before SW monsoon). The remaining months are mainly associated with rough weather, particularly from the end of June to August (SE monsoon).

**Table 1.** Summary per year of cetacean survey efforts (using all dataset) and cetacean sightings and numbers in Wakatobi National Park and adjacent waters.

| Year | No. of surveyed months* | Total no. of surveyed days | No. of days with sightings | No. of sighting events | No. of sighted individual cetaceans |
| --- | --- | --- | --- | --- | --- |
| 2004 | 6 (8) | 30 | 15 | 20 | 454 |
| 2005 | 9 (12) | 72 | 36 | 50 | 1,032 |
| 2006 | 7 (12) | 93 | 30 | 41 | 1,944 |
| 2007 | 12 (12) | 195 | 51 | 75 | 3,058 |
| 2008 | 12 (12) | 141 | 32 | 46 | 2,310 |
| 2009 | 6 (12) | 70 | 21 | 34 | 1,481 |
| 2010 | 8 (12) | 36 | 23 | 36 | 1,392 |
| 2011 | 5 (12) | 21 | 17 | 51 | 1,032 |
| 2012 | 3 (5) | 13 | 4 | 5 | 143 |
| Total | 68 (97) | 671 | 229 | 358 | 12,846 |

\*Number in parentheses indicates the number of complete months should be surveyed for that year. Only in 2007 and 2008, all months were surveyed. The study period lasted from May 2004 until May 2012.

**Figure 2.**
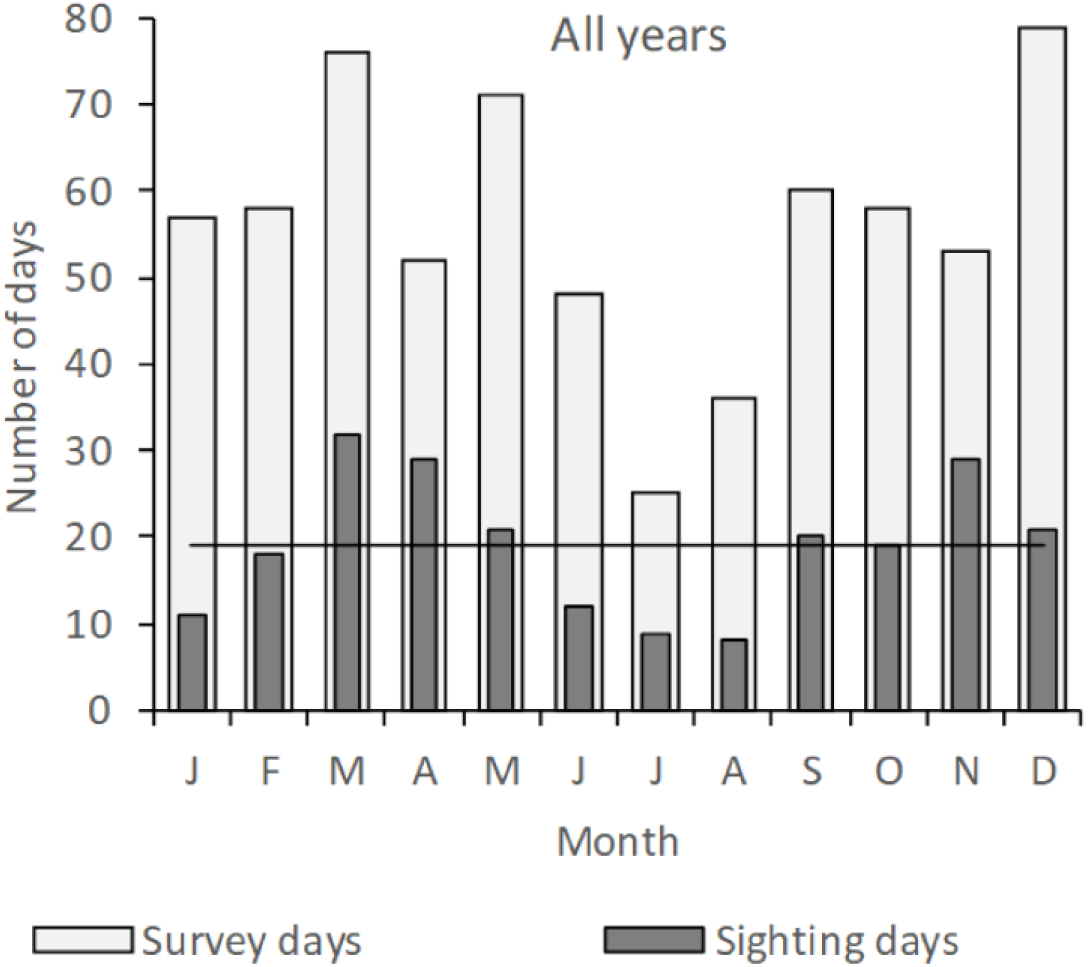
Total monthly survey efforts (number of survey days) and numbers of days with cetacean sighting during the Wakatobi cetacean monitoring program from May 2004 to May 2012. Horizontal line (—) indicates mean number of days with cetacean sighting over all months. T1 – Transition 1 season (Mar-May), SE-Mons – SE Monsoon season (Jun-Aug), T2 – Transition 2 season (Sep-Nov), SW-Mons – SW Monsoon season (Dec-Feb).

### 2.3.2. Cetacean diversity

Eleven species were positively identified in the WNP and adjacent waters during this study, which accounts for 37.4% of all sighted cetaceans. Among the identified species, eight were small cetaceans and three were large whales (sperm, Bryde’s, and blue whales) (Table 2). The four most frequently sighted species were spinner dolphin (17.6%), bottlenose dolphin (7.5%), sperm whale (5.6%), and melon-headed whale (3.6%) (Table 2). The sightings of these four species comprised 34.4% of the total number of sighting events. Each of six other species only represent less than 1% of the sightings, while the majority of the sightings (55.3%) were unidentified dolphins (Table 2).

**Table 2.** Summary of cetacean species sighted in Wakatobi and relative number of sightings and individuals (in relation to total n) from May 2004 to May 2012. IUCN (www.iucnredlist.org) status: DD, Data Deficient; VU, Vulnerable; NT, Near Threatened; LC, Least Concern; n.a, not available.

| Common name | Scientific name | IUCN Status | Sightings |  | Individuals |  |
| --- | --- | --- | --- | --- | --- | --- |
|  |  |  | n | % | n | % |
| Spinner dolphin | *Stenella longirostris* | DD | 63 | 17.6 | 3,244 | 25.3 |
| Bottlenose dolphin* | *Tursiops truncatus* | LC | 27 | 7.5 | 1,340 | 10.4 |
|  | *Tursiops aduncus* | DD |  |  |  |  |
| Sperm whale | *Physeter macrocephalus* | VU | 20 | 5.6 | 54 | 0.4 |
| Melon-headed whale | *Peponocephala electra* | LC | 13 | 3.6 | 890 | 6.9 |
| Pantropical spotted dolphin | *Stenella attenuata* | LC | 3 | 0.8 | 270 | 2.1 |
| Short-finned pilot whale | *Globicephala macrorhynchus* | DD | 3 | 0.8 | 140 | 1.1 |
| Cuvier's beaked whale | *Ziphius cavirostris* | LC | 2 | 0.6 | 3 | <0.1 |
| Risso's dolphin | *Grampus griseus* | LC | 1 | 0.3 | 5 | <0.1 |
| Bryde's whale | *Balaenoptera brydei* | n.a | 1 | 0.3 | 4 | <0.1 |
| Blue whale | *Balaenoptera musculus* | EN | 1 | 0.3 | 1 | <0.1 |
| Unidentified dolphins |  |  | 198 | 55.3 | 6,813 | 53.0 |
| Unidentified whales |  |  | 26 | 7.3 | 82 | 0.6 |
| All species |  |  | 358 | 100 | 12,846 | 100 |

\* During monitoring, the observers recognized two types of bottlenose dolphins: Common- (*Tursiops truncatus*) and Indo-Pacific bottlenose dolphin (*Tursiops aduncus*), however no clear difference in visual appearance of both species was evident and they could mostly not distinguish both species. Therefore, hereafter we refer to both species as bottlenose dolphins or *Tursiops spp.* whenever applicable.

The order of most sighted species based on the number of individuals was a bit different for the first four highest ranks: spinner dolphin (25.3%), bottlenose dolphin (10.4%), melon-headed whale (6.9%), and Pantropical spotted dolphin (2.1%). The greatest proportion of individuals reported were unidentified dolphins (53%). The identified cetacean species mostly have the IUCN status of ‘Least Concern’ (5 species) and ‘Data Deficient’ (3 species) (Table 2). Observations of less commonly sighted cetaceans also contribute to species presence information for endangered and vulnerable species in Wakatobi waters i.e. blue whale and sperm whale.

### 3.3. Spatial and temporal occurrence patterns

Overall, sightings were mainly concentrated in two geographic areas representing reef habitat types within the WNP i.e. inshore the main islands and south atolls, although some sightings were also observed in offshore outer reefs (Figures 4 and 5A). Spinner dolphins were seen in all habitat types (Figures 4A and 5A), while bottlenose dolphins and melon-headed whales occupied mainly the waters between the main islands and south atolls (Figures 4B,D and 5A). Sperm whales were sighted mostly in the east of the main island bordering outer reefs and in the north part of the main islands, and tended to avoid the south atolls (Figures 4C and 5A).

**Figure 4.**
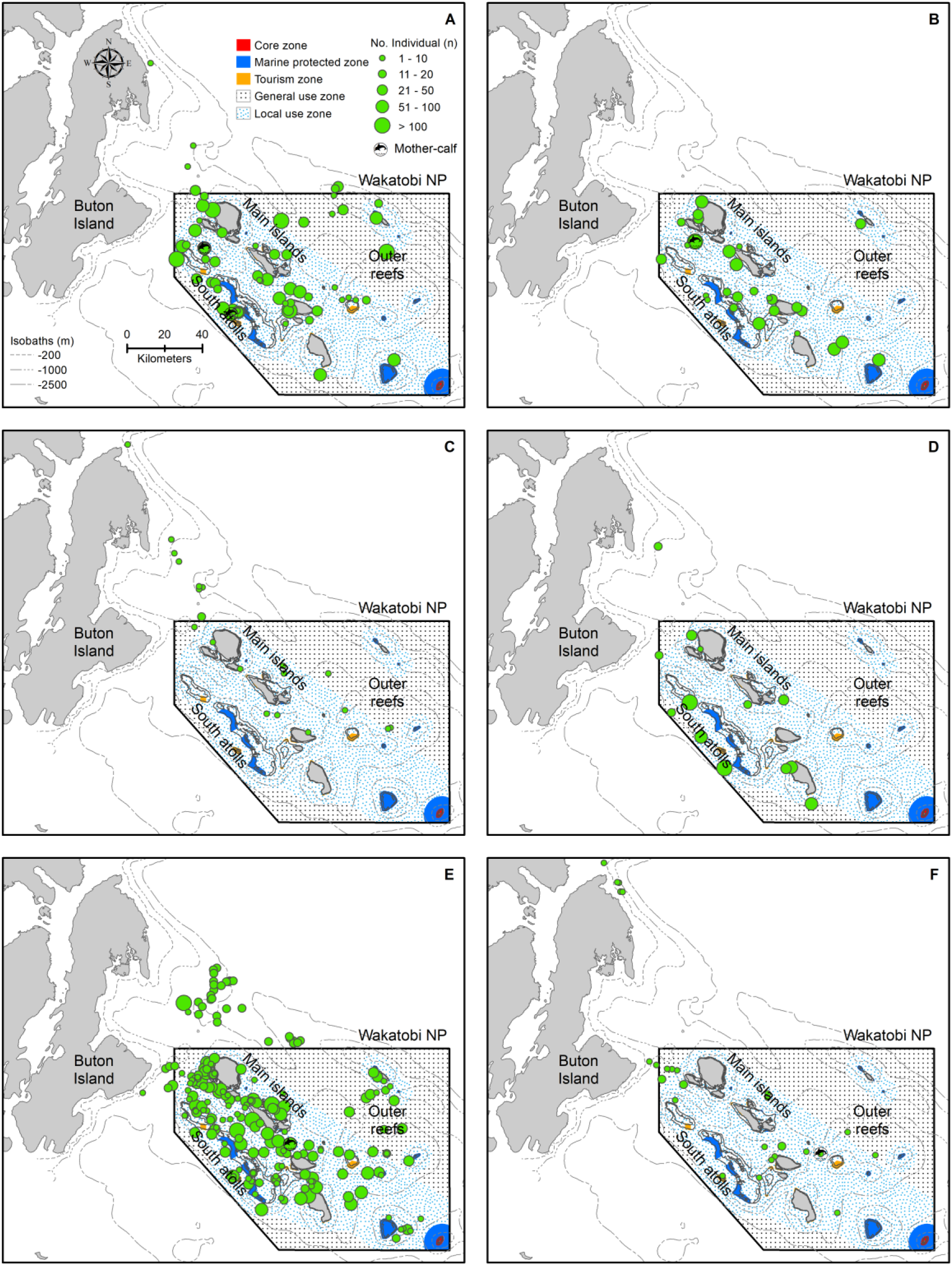
Spatial occurrence patterns of cetaceans in Wakatobi based on sightings in the period of May 2004 to May 2012: A. Bottlenose dolphin, B. Spinner dolphin, C. Sperm whale, D. Melon-headed whale, E. Unidentified dolphins, and F. Unidentified whales. The depth contours (isobaths) and Wakatobi National Park (WNP) zoning system are presented in panel A.

**Figure 5.**
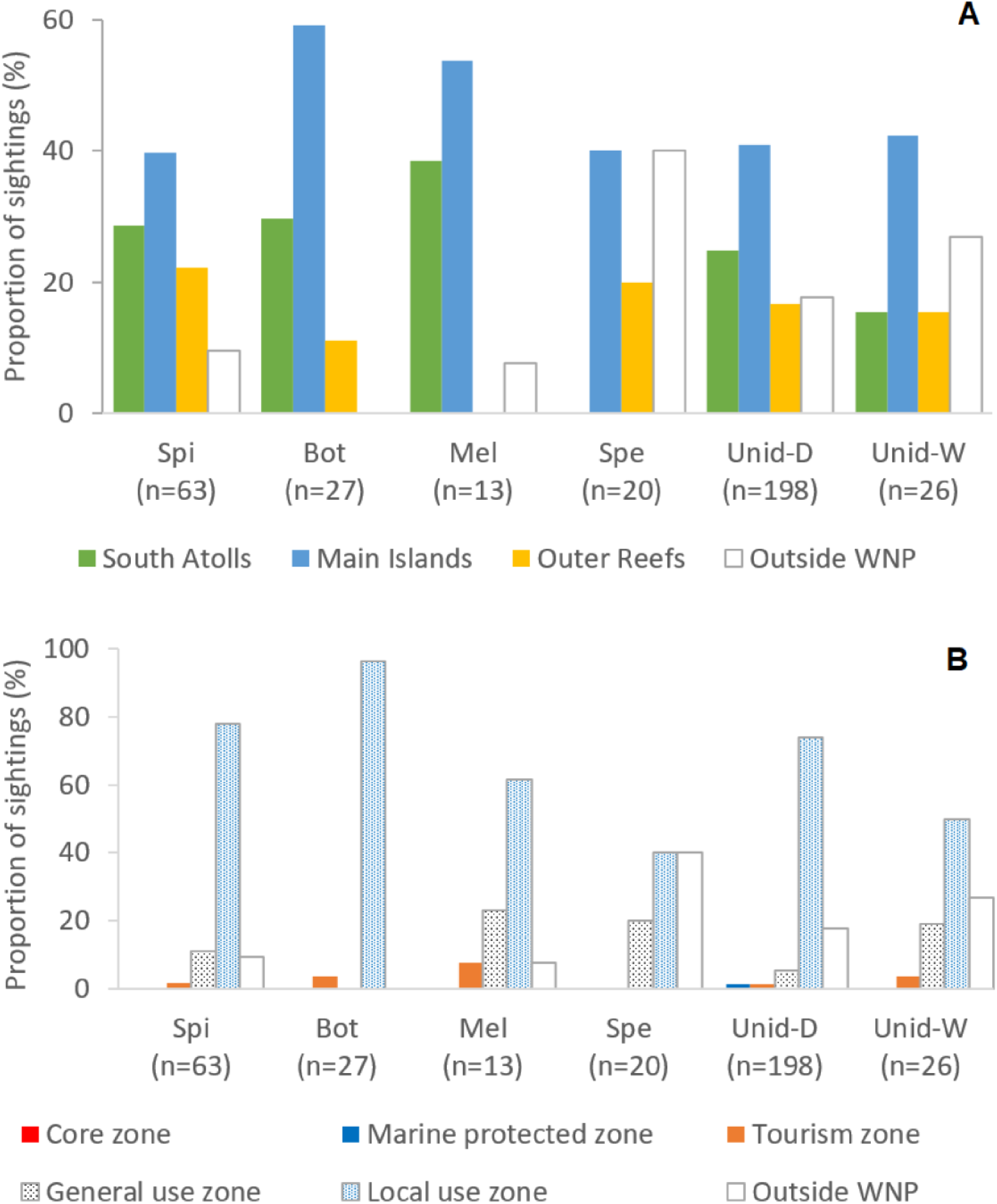
Proportion of sightings based on reef habitat types (A) and WNP zoning system (B). Bot-Bottlenose dolphin, Spi-Spinner dolphin, Mel-Melon-headed whale, Spe-Sperm whale, Unid-D – Unidentified dolphins, Unid-W – Unidentified whales.

Most of the cetacean sightings (>90% for small cetaceans, >60% for large cetaceans) occurred in the zones of the Wakatobi National Park that are also used by people. The WNP Authority accommodates three zones based on the types of human activities: tourism zone, local use zone (only available for local fishermen), and general use zone (open for all, including fishermen from outside WNP with permits) (WNP Authority, 2008), hereafter together indicated as ‘use zones’ (Figure 5B). The ‘no-use’ zones can be divided in a core zone (no go zone) and a marine protected zone (accessible but no human activities allowed). In addition to occurring in the use zones, sperm whale and unidentified whales were also prevalent outside the WNP zoning system (Figure 5 A,B). For all species and taxa, sightings seen within ‘no-use’ zones were less than 2%, while no sightings were even found in the core zone (Figure 5B).

Each of the four most sighted species had different depth preferences. Spinner dolphin and bottlenose dolphin mostly occupied shallower waters (depth median of −153m and −104m, respectively), while sperm whale and melon-headed whale were mostly seen in the deeper waters (depth median of −816m and −762m, respectively) (Figure 6), and sperm whale especially along the steep slopes (Figure 4C).

**Figure 6.**
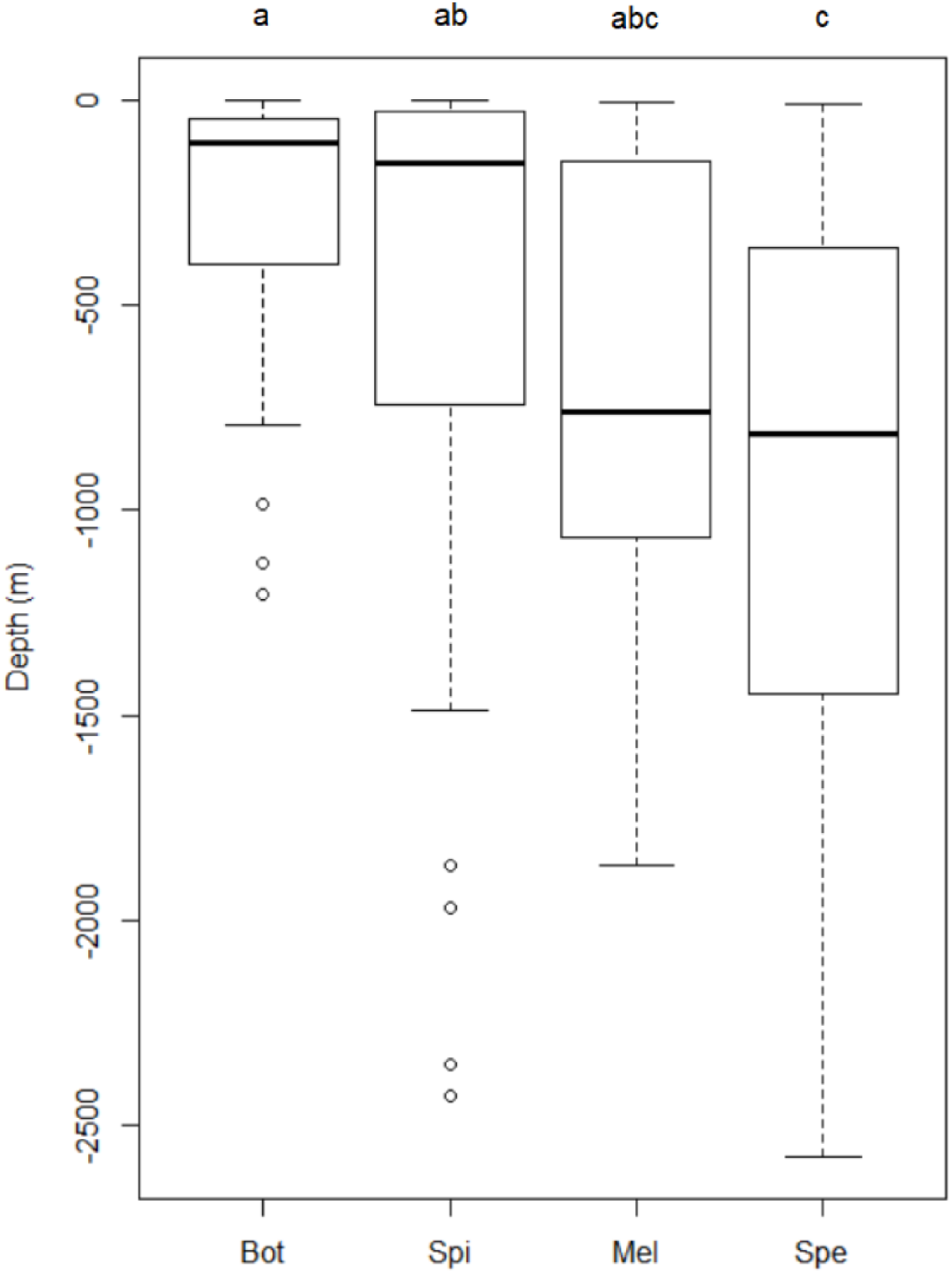
Boxplots of sightings of four most abundant cetacean species associated with depth in Wakatobi: Bot-Bottlenose dolphin (n=27), Spi-Spinner dolphin (n=63), Mel-Melon-headed whale (n=13), Spe-Sperm whale (n=20). Significant differences of depth among species were checked using Kruskal-Wallis. Wilcoxon post hoc test was applied for multiple comparisons after Kruskal-Wallis testing indicated a significant difference. The same letters indicate the boxplots do not differ significantly.

Seasonally, the number of identified cetacean species sighted was higher (>4 species) in March and April and from September to December, with a maximum of six species in November (Table 3). Only 2-3 identified species were recorded for the other months. Even though the data were collected through opportunistic survey platforms during all months, no successfully identified species was sighted in July (Table 3). Consequently, no single identified species was sighted year-round. Only in March, April, and November were the four most sighted species recorded (Table 3). No apparent temporal occurrence pattern was identified for the remaining six less sighted species.

**Table 3.** Seasonal occurrence: whale and dolphin species sighted in different months between May 2004 and May 2012 in Wakatobi (using all dataset). (+) indicates species was sighted at least once during the month.

| Species | Monsoon |  | T1 Season |  |  | SE-Monsoon |  |  | T2 Season |  |  | SW- |
| --- | --- | --- | --- | --- | --- | --- | --- | --- | --- | --- | --- | --- |
|  | Jan | Feb | Mar | Apr | May | Jun | Jul | Aug | Sep | Oct | Nov | Dec |
| Spinner dolphin | (+) | (+) | (+) | (+) | (+) |  |  | (+) | (+) | (+) | (+) | (+) |
| Bottlenose dolphin | (+) | (+) | (+) | (+) | (+) | (+) |  | (+) | (+) |  | (+) |  |
| Melon-headed whale |  |  | (+) | (+) |  | (+) |  | (+) | (+) |  | (+) | (+) |
| Sperm whale | (+) |  | (+) | (+) |  |  |  |  |  | (+) | (+) | (+) |
| Pantropical spotted dolphin |  |  |  |  |  |  |  |  | (+) | (+) |  |  |
| Short-finned pilot whale |  |  |  |  |  |  |  |  |  |  | (+) | (+) |
| Risso's dolphin |  |  |  |  | (+) |  |  |  |  |  |  |  |
| Bryde's whale |  |  |  |  |  |  |  |  |  |  |  | (+) |
| Cuvier's beaked whale |  |  |  |  |  |  |  |  |  |  | (+) |  |
| Blue whale |  |  |  |  |  |  |  |  |  | (+) |  |  |
| Number of delphinid species sighted | 2 | 2 | 3 | 3 | 3 | 2 | 0 | 3 | 4 | 2 | 5 | 3 |
| Number of large whales species sighted | 1 | 0 | 1 | 1 | 0 | 0 | 0 | 0 | 0 | 2 | 1 | 2 |
| All species sighted | 3 | 2 | 4 | 4 | 3 | 2 | 0 | 3 | 4 | 4 | 6 | 5 |

### 3.4. Behaviors, mother-calf pairs and cetacean-fishing vessel interaction

From 193 sightings (53.63%), information on behavior, mother-calf pairs or cetacean-fishing vessel interaction was recorded. Given that behaviors are species-specific, we showed the behavior per species for four most sighted cetaceans (Figure 7). Foraging and socializing were the two prominent behaviors in small delphinids (spinner dolphin, bottlenose dolphin, and melon-headed whale). Bow riding was prevalent in spinner and bottlenose dolphins. Only sperm whales were observed resting, although it is important to note that we only sighted three individuals for this species. For complete information on cetacean behaviors per species and taxa, see Table S1 in the Supplementary Information.

**Figure 7.**
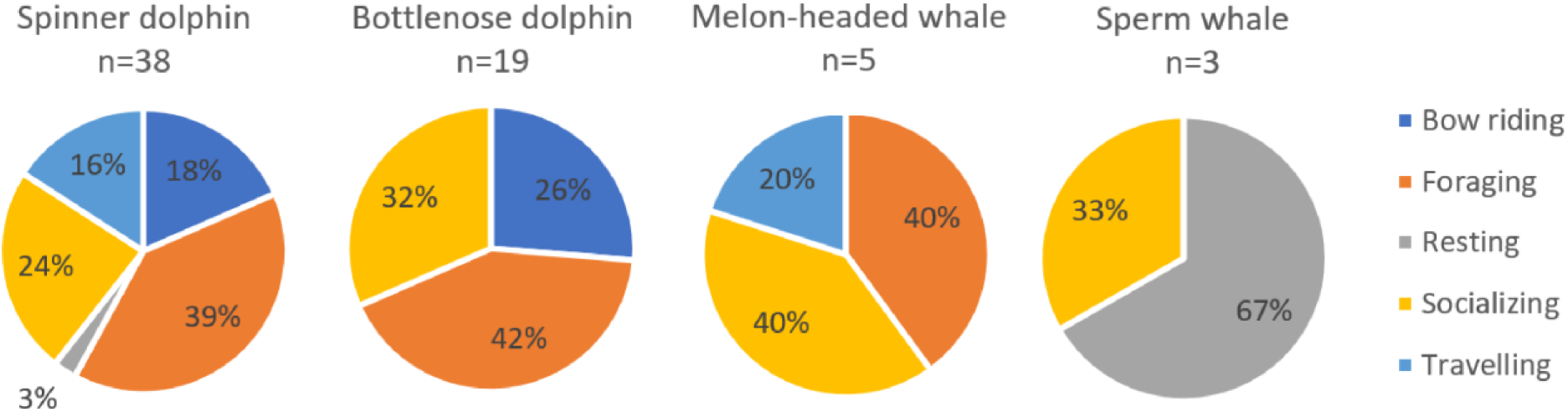
Relative occurrence of each behaviour category for four most abundant species.

Cetacean sightings with calves were observed five times (1.4% of total sightings) during the course of study: twice for spinner dolphin, and once each for bottlenose dolphin, unidentified dolphins, and unidentified whales. In total, 13.4% of the sightings were associated with fishing vessels or FADs; all were small cetaceans consisting mainly of unidentified dolphins and spinner dolphin, and to a lesser extent pantropical spotted dolphin, bottlenose dolphin and melon-headed whale.

### 3.5. Opportunistic survey results (sighting frequency)

A subset of all the surveyed days was defined as “eligible surveys” with comparable spatial coverage between months and years. The results showed that sighting frequency of large cetaceans was relatively constant, at around 0.25 sightings per day from 2004-2009, except for 2006 when it was <0.2 sightings per day. No sightings were recorded for large cetaceans in the last three years (Figure 3A). Small cetaceans were seen regularly over the study period with the sighting frequency ranging from 2.25 to 4.33 sightings per day, with three peaks occurring in 2005, 2010 and 2011 (Figure 3C). Monthly sighting frequency for both taxa showed a bimodal distribution, with peaks in September to November (Transition 1 season) and around March (Transition 2 season) (Figure 3B,D). The monthly trend was quite similar both for large and small cetaceans, i.e. months with no sightings for large cetaceans tend to also have less sighting frequency for small cetaceans (Figure 3B,D). The frequency of individuals per daily survey for both taxa can be seen in Figure S3 of the Supplementary Information.

**Figure 3.**
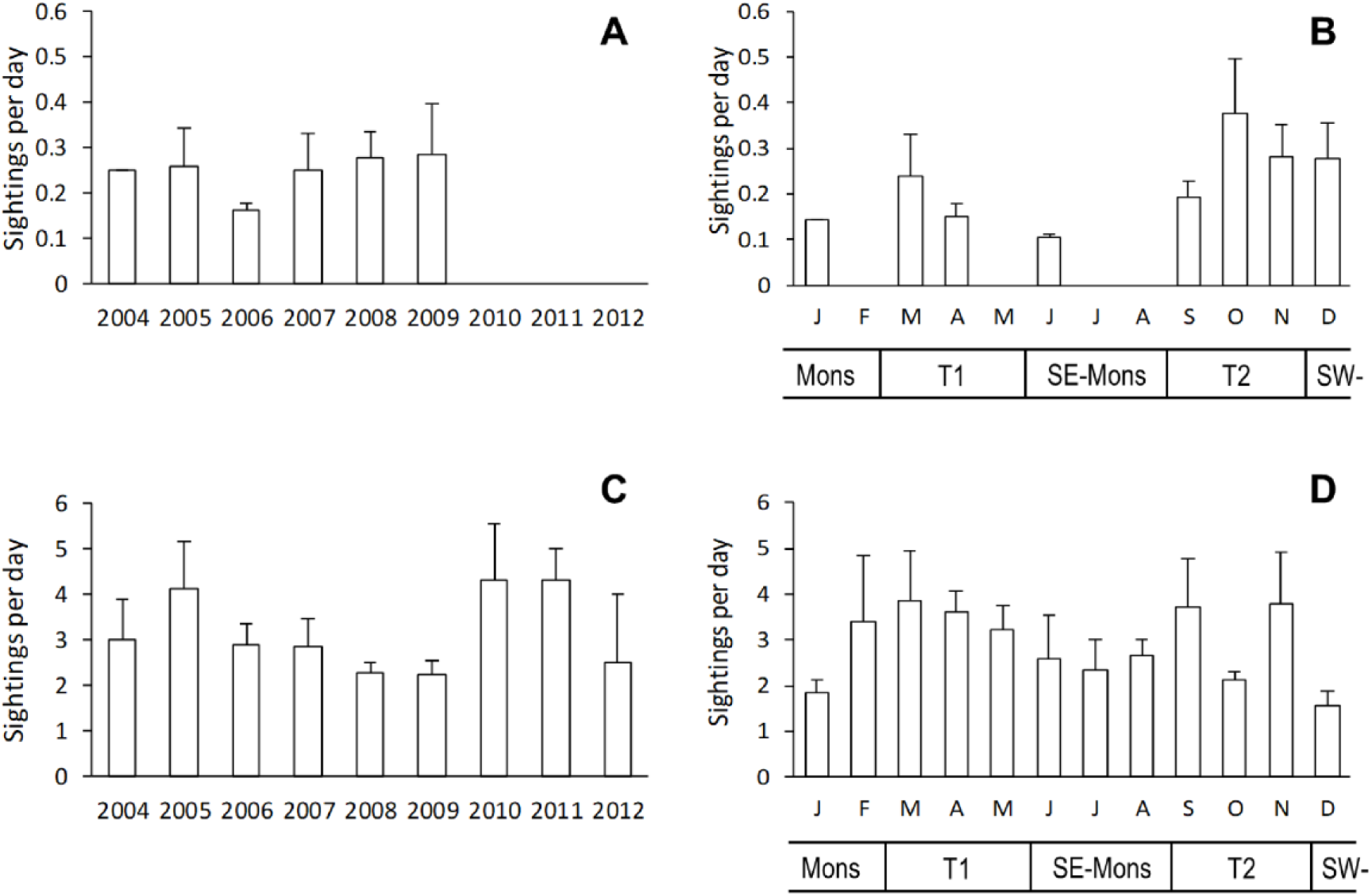
The average sighting frequency per day for each year from May 2004 to May 2012 for large cetaceans (A) and small cetaceans (C); and by month for large cetaceans (B) and small cetaceans (D). T1 – Transition 1 season (Mar-May), SE-Mons – SE Monsoon season (Jun-Aug), T2 – Transition 2 season (Sep-Nov), SW-Mons – SW Monsoon season (Dec-Feb). The sighting frequency were calculated from a sub-sample of dataset with surveyed days of 4 or more days per survey. Error bars indicate standard errors.

Calculating the number of all cetacean sightings per year and month allowed for a temporal analysis of cetacean occurrence over time (Figure S2, Supplementary). The number of sightings for large and small cetaceans fluctuated greatly over the years with the highest numbers in 2007 and 2008 for both taxa (Figure S2-A,C, Supplementary). Note that in 2004 and 2012, the survey efforts only cover parts of the year. The monthly sighting frequency (Figure 3B,D) is related to the monthly number of sightings (Figure S2-B,D, Supplementary). The total number of individual animals per year and per month for both taxa are presented in Figure S4 of the Supplementary Information.

### 3.6. Determinants to unidentified cetaceans

We conducted separate logistic regressions to examine possible determinants to unidentified dolphins (n=198) and unidentified whales (n=26), using season, distance between the observers and the animal group size as the independent variables. Season and distance were not significantly associated with the prevalence of unidentified dolphins or whales. The number of dolphins per group per sighting was a significant determinant for the prevalence of unidentified dolphins (p=0.000, B= −0.240, Exp(B) or ratio = 0.787), which was produced by a model with Block Chi Square =56.650 (p model=0.000, Nagelkerke Rsqr=0.364). Using the same model setting, the number of whales per group per sighting was also a significant determinant for the prevalence of unidentified whales (p=0.000, B= −0.240, Exp(B) or ratio = 0.787, Block Chi Square =56.650, p model=0.000, Nagelkerke Rsqr=0.364). Thus, the smaller the number of animals per group was, the more likely it was that the species was unidentified (possibly because it was harder to identify the animal with only a few individuals being present).

## 4. DISCUSSION

This is the first published study of cetaceans in Wakatobi National Park and adjacent waters. The data was collected mainly using platforms of opportunity in the period of May 2004-May 2012. It presented a unique opportunity to document the occurrence and habitat use of yet unstudied island-dependent oceanic cetacean populations. Here, we provide new information on what cetacean species occur around this archipelago and how their presence changes over time. In addition, sightings provided information on the presence of mother-calf pairs, cetacean behavior and interactions with fishing vessels and FADs. We also discuss the implications of our findings for the conservation of these large marine top predators and for marine park management.

### 4.1. Caveats

A number of caveats are inherent when using platforms of opportunity. Some issues can be addressed during the analysis stage; others will lead to limitations in how the data can be interpreted.

Design-based survey transects are planned to ensure that every point in the study area has the same chance of being surveyed (Buckland et al., 2001). Platforms of opportunities like ferries can follow regular routes, but those routes do not provide a representative coverage of cetacean habitats. There are some methodological approaches, such as spatial modelling, where this type of data can be used to obtain distribution maps in areas that were covered poorly (Hedley and Buckland, 2004). This approach was not possible during this study due to the logistical and data quality constraints we will discuss further below. Another approach that has been used is to analyze presence-only data to obtain more complex habitat suitability models for cetacean species or taxa (Kaschner et al., 2006).

The survey data was collected under different conditions that influence the sighting probability for cetaceans. This includes vessel characteristics, observation techniques, observer experience as well as environmental conditions. For example, the observation height varied among platforms (0.5–5m) as well as vessel speeds (7-20 knots). The number of observers varied (at least one person), observers used different sighting methods (unaided eyes, binoculars) and they had different levels of experience in species identification and group size estimation. Weather conditions were not recorded, although vessels did not go out in sea states of Beaufort >4, making the sighting conditions relatively comparable between surveys. In addition, cetacean species have differing chances of being visible, for example due to their dive behavior (Barlow, 2015). Ideally the monitoring results are corrected for such factors (Evans and Hammond, 2004). Only a simple sighting frequency per surveyed day was calculated in our study, likely leading to an unquantified bias.

Effort differed between survey areas and over seasons, meaning that the spatial effort could only be approximated. This circumstance makes it difficult to ascertain if the survey coverage of the study area by the different platforms of opportunities was similar between months or years. This event means that there is a chance that the observed spatial patterns may reflect the distribution and abundance of the vessel effort rather than of the animals.

Due to the lack of GPS tracks, sighting frequency could not be reported as the number of sightings per km or per hours surveyed, as is commonly done (Alves et al., 2018). Instead effort was recorded as days surveyed. This fact has been done for other studies using platforms of opportunity (e.g. Pérez-Vallazza et al., 2008) and allows the sighting frequency to be determined, reported as ‘sightings per day’. Assuming that sighting records are obtained throughout the study area, the results can still provide a good qualitative indication of the distribution and abundance of animals (Berrow et al., 1996). In our study, this assumption could be done for a subset of data, i.e. mainly the fishery monitoring surveys, where more consistent information on spatial coverage was available. Some of the limitations of this data set are discussed further below.

The value of opportunistic surveys in the WNP and surrounding waters could be improved by including GPS tracks, which are much more affordable nowadays considering recent technological developments. Also, the use of photographs of observed animals could help with species identification and greatly contribute to the value of opportunistic monitoring.

Still, even considering these caveats, the data collected during the non-systematic surveys enable the construction of a positively identified species list and a first indication of the cetacean population status and density. The applied method provides a low-cost approach to obtain valuable long-term information for areas with limited or no specific monitoring effort over a wide geographic area (Delefosse et al., 2018). Although interpretation of non-systematically collected data is difficult without measure of effort (Evans and Hammond, 2004), with stringent data filtering and quality control, valuable information can be obtained (Pikesley et al., 2012).

### 4.2. Cetacean diversity

Cetaceans are an important component of marine ecosystems as apex predators that influence structure and function of marine communities (Foley et al., 2010). Information on cetacean species around Indonesia was primarily based on strandings (Branch et al., 2007; Mustika et al., 2009) and incidental sightings (Rudolph et al., 1997). Only recently, cetacean monitoring programs have been conducted in several sites, including Papua, East Kalimantan and wider Lesser Sunda (Bali, Savu Sea, Alor) (Ender et al., 2014; Kreb, 2004; Mustika, 2006). For Wakatobi waters the limited multi-species studies on this taxonomic group that have previously been performed are only available in grey literature. This study is the first to provide important baseline information on the ecology and the community patterns of cetaceans in Wakatobi waters. Eleven cetaceans were positively identified at species level representing 32% of all cetacean species recorded in Indonesia (Mustika et al., 2015b). These include species featured in the International Union for Conservation of Nature (IUCN) Red List of Threatened Species (IUCN, 2020) as Endangered (blue whale, *Balaenoptera musculus*), and Vulnerable (sperm whale, *Physeter macrocephalus)*. Almost two thirds of the identified species are listed as Least Concern or Data Deficient, the last category making it difficult to assess their population status. WNP has a high habitat biodiversity which is reflected in the reported species that occupy different habitats. This finding highlights the important role of Wakatobi waters as a regional reservoir of cetacean diversity. Throughout the study period (from 2004 to 2012) 90% of the species had been identified within the first 5 years. This result indicates that even with the non-systematic coverage, the most prevalent and possibly resident species were detected early in the study. Nevertheless, it is important to note that the majority of sightings (62.6%) were recorded as unidentified dolphin or whale species, so there are probably still a number of undetected species. Even during dedicated surveys, identification rates for small cetaceans can be below 50% (Di Tullio et al., 2016). Therefore, species identification training for local observers and taking photos during the sightings are imperative to increase identification success rate in the future. Taking photos are particularly important to increase the likelihood of identifying animals in small groups or single animal.

### 4.3. Sighting frequency

Assuming that survey effort is roughly similar between surveys, the sighting frequencies can provide a long-term measure of occurrence and changes for a study area. These results tend to be more quantifiable than studies that only use incidental sightings with no associated effort data (Evans and Hammond, 2004), or are based on perceived trends of cetacean abundance reported by fishermen (e.g. Maynou et al., 2011).

In this study, sighting frequencies were used for a subset of the data which allowed the investigation of temporal changes in cetacean occurrence between seasons and years for Wakatobi waters. The reduced number of records made it necessary to pool all sightings into either the category small or large cetacean. The sighting frequency for both small and large cetaceans, combined for the years 2004 to 2012, indicated a bimodal distribution with peaks in the two transition seasons, September to November and March. There are a number of possible reasons for the observed patterns. Weather conditions are calmer during the inter-monsoonal season than in the generally much rougher SE and SW monsoons (Pet-Soede and Erdmann, 2003). As discussed earlier, data on weather conditions, such as sea swell or wind strength, were not recorded during the survey. The vessels generally only went out at sea to around sea states of Beaufort < 4, which makes the weather conditions relatively comparable between surveys. For species that occur in large groups, approach vessels for bow riding and show active surface behavior, such as spinner dolphins, the sighting probability would likely not change within this range. However, group size plays an important part in detectability. For smaller groups or animals or those that are only visible for short times, the chance that an observer sees them can be drastically reduced, for example when white-caps occur (Beaufort 3) (Barlow, 2015). This is in particular true for cryptic species such as beaked whales that occur in solitude or in small group sizes and are difficult to spot (and identify) even under good sighting conditions.

Another challenge when interpreting the observed seasonal patterns of density is the pooling of all species into either the category small or large cetaceans. This challenge makes it impossible to see in what way changes in distribution of different species within each category, e.g. due to changes in prey availability, might influence the overall pattern. This circumstance highlights again the importance of having a higher number of identified sightings which would allow for the investigation of species specific changes in occurrence, both throughout the year and interannually.

For this analysis we considered a survey effort of at least four days to be the minimum requirement to be used for the calculation of a sighting frequency. This arrangement however did not mean that survey effort was the same over time. Due to limited resources at the end of the monitoring program, survey effort was lower in the last three monitoring years (Figure S1, Supplementary). During those three years no large cetaceans were sighted (Figure 3A). This is most likely because the offshore area received less survey coverage, especially in the outer reefs where large cetaceans were seen more often than in main islands and south atolls (A. Sahri and Purwanto, pers. obs.). Conversely, with the same survey effort around the much smaller inshore waters, the small cetacean density has increased in the last three years (Figure 3C). It is possible that for species that are common, such as spinner and bottlenose dolphins, the sighting frequency is quite robust as they are also seen frequently in the times of reduced effort. For species sighted more rarely, such as large whales, a reduced effort can lead to no more animals being registered, thus making the calculation of sighting frequency impossible.

Without further study though, it cannot be excluded that a decrease in number of sightings could also be related to depletion of the populations (Amir et al., 2012) or a change in distribution. It is important to use similar temporal and spatial coverage between years, even when using platforms of opportunity. The availability of abundance indices such as sightings per km or hour (rather than sightings per day, as used here) would also provide more refined insights into seasonality of cetacean presence in the study area.

### 4.4. Spatio-temporal occurrence patterns

This study was able to provide a preliminary overview of the spatial distribution of cetaceans in Wakatobi over almost a decade. As records used in this study are based on non-systematic sightings, the absence of standardized design make it impossible to evaluate the absolute species abundance (Robinson et al., 2013).

The complex Wakatobi ecosystem can attract and support a variety of cetacean species, because Wakatobi is located in the Indonesian tropical upwelling system, where oceanic currents strongly stimulate primary productivity (Drushka et al., 2010; Steinke et al., 2014). Species found here range from shallow water species to offshore pelagic species. Spinner dolphin, bottlenose dolphin, and melon-headed whale were observed in the vicinity of fringing reefs around the coastline, atolls and small islands. Sperm whales were abundant in the vicinity of the submarine trough-like features between major landmasses (Pet-Soede and Erdmann, 2003) in the east part of the main islands bordering the outer reefs, and in the deep waters in the north of the main islands. The numbers of sightings for the other six species were too low to present reliable occurrence patterns, but they are included here for completeness as a first indication of cetaceans present within this region. Additional effort is necessary to obtain more information about these rare and cryptic species in order to identify areas that are most important for conservation and management actions. The co-occurrence of different species in Wakatobi waters offers a great opportunity to investigate possible interactions among species as well as their habitat selection on a local scale (Dellabianca et al., 2018). This area is unique as both coastal and oceanic cetacean species can occur in WNP due to the presence of deep water very close to the coast. This feature allows the investigation of species that normally occur in deep waters further offshore, which in other areas are too challenging to monitor with small boats (Ponnampalam, 2012).

The narrow and very steep continental shelf provides deep waters close to the shore, and the presence of submarine features around the coastline probably contributes to upwelling, thus enhancing the productivity of the waters that support the feeding requirements of several species distributed around the islands. This fact has been shown before in other parts of the world (Bouchet et al., 2015; Rennie et al., 2009). Oceanographic processes such as the monsoonal regime with seasonally reversed currents lead to water exchange between the Flores and Banda Seas and drive the seasonal upwelling in these waters (Pet-Soede and Erdmann, 2003). This condition may attract cetaceans through enhanced productivity and related prey availability (Baird et al., 2009).

Some spinner dolphin populations are island-associated and known to move between coastal and oceanic habitats (Dolar et al., 2006; Ponnampalam, 2012). In Hawaiian waters, such regular diel movements have been linked to prey migrations (Benoit-Bird and Au, 2003). For the Sulu Sea, spinner dolphins were found to primarily feed on mesopelagic fishes in depths of 200-400m as well as squid and crustaceans (Dolar et al., 2003). Surveys in late November and early December around the Raja Ampat archipelago found spinner dolphins to be the most commonly sighted cetacean, occurring in shallow shelf as well as deep oceanic waters (Borsa and Nugroho, 2010). The results presented here for Wakatobi waters are in line with what has been recorded for spinner dolphin populations in other island and archipelago habitats.

A similar distribution pattern was seen for bottlenose dolphins. The observed inshore bottlenose dolphins may have been *T. aduncus*, while those observed offshore may have been *T. tursiops*. These species are very difficult to distinguish at sea, in particular if they are not approached and/or photographed. Consequently, both species were recorded as bottlenose dolphin *(Tursiops* spp). Bottlenose dolphins are known to form stable social groups, sometimes with small resident populations that have a high affiliation to a relatively small area (Dulau et al., 2017; Passadore et al., 2018a; Shirakihara et al., 2002). These populations seem to be linked to coastal areas that have high prey availability and low predation risk (Haughey et al., 2020). The distribution data from Wakatobi indicates that bottlenose dolphins are occurring wide-spread in the area on a regular basis, giving rise to the possibility that at least some of these animals are staying in these waters continuously. World-wide bottlenose dolphin populations can be genetically distinct at small spatial scales, which is highly relevant for any conservation efforts (e.g. Chen et al., 2017). The use of photographs of the dorsal fin of individual animals, as well as sampling of tissue for genetic analysis, could provide more insight into this.

Sperm whale and unidentified large whales were seen significantly more outside the WNP and in deep waters to the north of the main islands. Sperm whales were mostly reported in deep water which matches the feeding trait of the species; they feed mainly on mesopelagic squid and fish around the shelf edge (Whitehead, 2009). The local depth and seabed configuration (Pikesley et al., 2012) in combination with oceanographic processes (Tynan et al., 2005) are associated with aggregated cetacean prey species (Bearzi et al., 2008). Baleen whale sightings were comparatively rare. In contrast to other baleen whales, Bryde’s whales do not conduct long-distance migrations, instead remaining in tropical and warm temperate waters all year (Kato and Perrin, 2009). They likely conduct movements to areas where breeding occurs, however, the general knowledge on the behavior of this species is still poor (Kato and Perrin, 2009). Sightings around Wakatabi could indicate animals using this area on a regular basis, but further data is needed to confirm this.

Several species were found to use Wakatobi waters on a regular basis, with the four most abundant species being present from 50% to 83% of the year (Table 3). Only in July, each of three sightings of unidentified dolphins occurred in 2006, 2007, and 2010, much less than other months. The most reasonable explanation is that this is related to the very rough Wakatobi waters in this month in the middle of the SE monsoon limiting survey efforts (Figures 2 and S1). While one would in general assume that the productivity in the area was high during the summer, it could also be that this was not the case. The productivity of the waters around the Indonesian Archipelago, and in particular the Banda Sea, are influenced by large-scale physical processes as well as local factors such as surface wind and rain (Moore et al., 2003). There are a number of complex interactions, and in addition human activities on land, leading to a high input of nutrients, and the change in temperature due to climate change, leading to a more stable stratified layer and less upwelling (Chang et al., 2019). Monitoring of the physical processes, productivity as well as the occurrence of top predators would help to achieve a better understanding of the relationship between these factors in Wakatobi waters. Consideration of cetacean monitoring technologies that can also function well during the rough season, such as stationary passive acoustic devices, can clarify this issue (Anderson et al., 2012a; André et al., 2011).

### 4.5. Behaviors, mother-calf pair presence, and cetacean-fishing vessel interaction

This study also shows that all vital activities of cetacean life cycle such as foraging, socializing, resting, travelling (Figure 7), and calving occurred in Wakatobi waters. The foraging strategy in small cetaceans such as spinner and bottlenose dolphins was associated with depth since these animals usually follow the vertical movements of their prey (Benoit-Bird and Au, 2003; Torres and Read, 2009). Thus, information on cetacean foraging depth is important for ecological investigation. In Wakatobi, spinner and bottlenose dolphins forage in depth around −230m and −60m, respectively, reflecting different depth preferences probably to avoid prey competition. Out of three sperm whales with recorded behaviors in our dataset, two of them were resting. Resting behavior in large whales such as the sperm whales (Amano and Yoshioka, 2003; Miller et al., 2008) needs to be considered in the WNP spatial planning due to the possible risk of collision (Frantzis et al., 2019) with ships that are using the same area. In addition to the aforementioned need for increased skills for species identification, the identification of species behavior is also needed for future data collection. The presence of some cetacean mother-calf pairs suggest that the WNP provides good nursery and calving grounds for several cetacean species.

Some of the small cetacean species are gregarious by nature. The largest group of cetaceans sighted during these surveys comprised of an estimated 200 individuals of spinner dolphins, in line with other studies reporting large groups of up to 500 (De Vos et al., 2012) or even 1000 individuals (Gore et al., 2012). Spinner dolphins are also known to show impressive surface behavior, such as their “spinning” (Würsig and Whitehead, 2009), making them highly visible. Two other species that were seen in groups of more than 100 individuals are melon-headed whale and bottlenose dolphin. Both spinner and bottlenose dolphins frequently approached vessels for bow riding. These charismatic and abundant species could potentially be the basis for ecotourism activities. Cetacean-watching can have great value if conducted in a responsible manner to avoid negative impacts on the animals (Mustika et al., 2015a). Generating economic value for local inhabitants will help to motivate them to protect the marine communities.

During the surveys, skipjack tuna *(Katsuwonus pelamis*) were spotted at the surface exclusively around fishing vessels and fish aggregating devices (FADs), subsequently attracting small cetaceans (A. Sahri, pers. obs.). Dolphins tend to aggregate in such areas where their main prey occurs to increase their feeding success rate (López et al., 2004). Fishermen in Wakatobi even use the presence of small cetaceans (mainly spinner and bottlenose dolphins) to locate the schools of yellowfin tuna (A. Sahri, pers. obs.) as has been described for other areas (Anderson, 2005; Anderson et al., 2012b). The co-occurrence between cetaceans and fishing activities increases the risk for cetacean entanglement and drowning (Whitty, 2015). Reeves et al. (2003) suggested that local declines in small cetaceans are mostly due to increased intensification of fisheries and vessel activities. In addition, the co-occurrence with fishing activities may cause fish stock depletion for the cetaceans (Gore et al., 2012).

### 4.6. Implications for marine park management

Knowledge of cetacean population presence and ecology is fundamental for formulating conservation policy (Reid et al., 2003) and an effective policy depends on understanding relationships between species, habitats and anthropogenic interactions (Cañadas and Hammond, 2008). The management of protected areas that is designed for top predators as umbrella species is highly efficient, resulting in higher biodiversity and more ecosystem benefits (Sergio et al., 2008). For wildlife conservation management, baseline knowledge is needed of the presence of cetacean populations and identification of areas of particular importance for specific species (Panigada et al., 2008). The approach and findings described here can therefore support the WNP Authority. Our results indicate that the main islands and south atolls (Figure 1) constitute areas of special interest for cetacean conservation as several species, especially small cetaceans, aggregate here and display different important behaviors including resting, socializing, foraging. The areas are also important for post-breeding activities such as nursing and calving. However, this area also coincides with important fishing grounds and tourism activities (Figures 4 and 5). Cetaceans in Indonesian waters have been under strict protection by law since the 1970s (Sahri et al., in press). All cetacean species in Indonesia are listed as protected animals in the annex of the Government Regulation No. 7/1999 (The Government of The Republic of Indonesia, 1999). Because of concerns about fishing and tourism activities on cetacean populations in protected areas (Howes et al., 2012; Read, 2008), it is of major importance to include this information in governmental management plans.

One of the most important approaches to marine conservation is the establishment of marine protected areas (MPAs) (Cañadas et al., 2005), although their effectiveness greatly depends on the level of enforcement. Wakatobi National Park was established with multiple-use zoning system, including two no-use zones (core and marine protected zones) and three use zones (tourism, local use, general use zones) (WNP Authority, 2008). Only ~3% of the total WNP area falls under the no-use zone, and this mainly concerns more sedentary coastal and marine ecosystems such as coral reefs, seagrass beds, mangrove, fish spawning aggregation sites and turtle and seabird nesting sites (WNP Authority, 2008). It is not surprising that most sightings for all cetacean species and taxa occurred within use zones, which is ~97% of the total area. No sightings were found in the core zone of the WNP, which is a crucial finding that indicates the mismatch between WNP design and the ecological needs of the cetaceans. Mobile species such as the cetaceans are hardly accommodated in the current WNP zoning system, because >90% of small cetacean sightings and >60% of large cetacean sightings occurred in the use zones in which traditional fishing and recreational activities are allowed to take place. These areas are commonly highly used corridors by both cetaceans and humans, implicating a potential conflict area. The adverse impact of fishing, shipping and tourism on cetaceans can be addressed by setting and enforcing stricter rules such as only applying cetacean-friendly fishing gear, restricting vessel speeds and developing responsible ecotourism. The WNP Authority has set rules on fishing gear that is allowed to be used within the WNP (WNP Authority, 2008), however no such a ‘code of conduct’ is available yet for whale and dolphin watching tourism, even under the national legislation (Sahri et al., in press).

Large whales, especially sperm whales, occurred in the use zones as well as outside the WNP boundaries. The latter areas do not have any regulations in place to manage potential threats to cetaceans. MPAs that are predominantly designed for protection of coastal habitats and to support sustainable local fisheries may have little effect in protecting highly mobile cetaceans (Dinis et al., 2016). Designing protected areas for specific species requires knowledge of the spatio-temporal distribution and habitat requirements of the targeted populations, in order to adjust the size of the management area to their ecological needs (Silva et al., 2012). To provide ample protection for the cetaceans, an adequate management regime (Howes et al., 2012) and MPA range expansion are needed, since the mobile animals usually have ranges that go outside of a single MPA (Dinis et al., 2016; Wilson et al., 2004). An MPA network may be needed to truly protect these highly mobile species (Hooker et al., 2011).

Our approach can be seen as an inexpensive pilot study to identify areas of high sightings that could inform future survey designs; identify candidate time-area fisheries closures e.g. to minimize bycatch; or direct ecotourism development, since the area demonstrates the potential for ecotourism based on whale and dolphin watching activities. The possible connectivity between the Wakatobi cetaceans and those of the Flores and Banda Seas need further investigation to strengthen the cetacean conservation in the central and eastern waters of Indonesia. These areas have recently been recognized to be important for conservation by an ongoing international research and conservation initiative, named Important Marine Mammal Areas (IMMAs) (IUCN-MMPATF, 2019). This recognition highlights the need for protecting this unique and vulnerable area through the adoption of substantial management measures informed by scientific evidence.

Information obtained from this study can help inform national and local governments with data for cetacean species with different status from Data Deficient to Endangered under the IUCN Red List of Threatened Species (IUCN, 2020). Distribution and species diversity data, for instance, are recognized as critical for managing cetaceans in Indonesia (Ender et al., 2014). The information is useful for driving further action on the conservation of the species. Information on behavior and their spatial and temporal occurrence of cetacean species is urgently needed in the WNP to inform the future management of specific threats to cetaceans and to identify appropriate areas for MPAs (Cañadas et al., 2005; Gómez De Segura et al., 2006). When cetaceans frequently use an MPA, cetacean conservation issues need to be included in the MPA management plan (Cañadas and Hammond, 2008; Dulau-Drouot et al., 2008). Information on cetacean ecology should be used to adjust the existing zoning system as an important step in balancing the needs of local users and those of protected species. Adjusting zoning designation of areas where threatening human activities significantly overlap with important cetacean habitat can contribute effectively to the species’ conservation (Silva et al., 2012).

Our study indicates a high species diversity of cetaceans in the poorly studied Wakatobi waters and provides a preliminary overview of cetacean spatial and temporal occurrence as well as their behavior and habitat preference. The results show that the presence of deep pelagic waters close to the islands makes it a unique habitat for not only coastal species, but also those that normally occur offshore and are much less accessible. Our objective was not so much to quantitatively assess cetacean abundance, but rather to evaluate whether non-systematic low-cost survey data can be a valuable information source in data-poor situations and where funding is limited. Relative occurrence, such as a sighting rate, if collected appropriately, is an adequate method for long-term monitoring and can provide the information needed to support improved park management and conservation planning. Some of the caveats that were identified during data collection can be easily addressed, such as the improvement of effort data by recording of GPS tracks. Data analysis and inferences based on non-systematic surveys need to take the limitations of the method into account. Where cetacean studies or wildlife-based ecotourism are just beginning, non-systematic surveys and collection of opportunistic sightings can provide enough information on animal occurrence that it can inform the kind of future studies that might be needed. We recommend that data collected from non-systematic, opportunistic, or incidental based projects be published to strengthen the value of volunteer efforts (Theobald et al., 2015) and to maximize the benefit of such efforts for science and management (Pirotta et al., 2019).

With some improvement to the current data collection protocols, more accurate estimates of cetacean density could be obtained. These would involve conducting appropriately randomized and stratified surveys and collecting and incorporating sighting parameters in the estimation of sighting rate. Using platforms of opportunity to assess cetacean occurrence is a low cost method. The consistent use of vessels that have another work focus, for example, is feasible as long as the effort is recorded. Continuous monitoring over several years can help assess impacts of larger scale changes in habitats, climate change and anthropogenic uses. More in-depth studies using systematic surveys are needed to provide more robust information. Localized studies of cetacean species focusing on residence patterns, genetic structure, and population impacts arising from interactions with other anthropogenic threats, such as marine traffic, should yield additional information for management strategies at the local level. Finally, we advise to consider protecting currently unprotected cetacean key-habitats, and setting stricter rules for human activities in the current use zones during future WNP rezoning processes.

## DATA AVAILABILITY STATEMENT

The data analyzed in this study is subject to the following licenses/restrictions: The data was obtained through a data sharing agreement from The Nature Conservancy (TNC) Indonesia, and therefore is not publicly accessible. Requests to access these datasets should be directed to Yusuf Fajariyanto.

## AUTHOR CONTRIBUTIONS

AS, PM, AM and MS conceived the ideas; AS and PP collected the data; AS, MS and PM analyzed the data; AS, AM and MS led the writing; AS, PM, PP, AM and MS discussed the results and commented on the manuscript.

## ACKNOWLEDGEMENTS

The cetacean data was collected by the Wakatobi National Park Authority and the Joint Program of The Nature Conservancy (TNC) Indonesia and WWF Indonesia. Authors would like to thank all Wakatobi National Park staff, Wakatobi District Government staff, speedboat captains and crews, monitoring staff, all volunteers who collected data in the fields and especially Yusuf Fajariyanto for data sharing agreement. Two authors AS and PP were employed as monitoring coordinators by TNC Indonesia to conduct data collection. The authors would like to thank the Indonesia Endowment Fund for Education (LPDP) Scholarship from Ministry of Finance of Indonesia who financially supporting this study. The authors would like to thank Debby Barbé for the preliminary study and the reviewers who helped to improve the quality of the manuscript.

## Footnotes

1 https://www.gebco.net/

